# Calcium channel-dependent induction of long-term synaptic plasticity at excitatory Golgi cell synapses of cerebellum

**DOI:** 10.1101/2019.12.19.882944

**Authors:** F. Locatelli, T. Soda, I. Montagna, S. Tritto, L. Botta, F. Prestori, E. D’Angelo

## Abstract

The Golgi cells, together with granule cells and mossy fibers, form a neuronal microcircuit regulating information transfer at the cerebellum input stage. Despite theoretical predictions, little was known about long-term synaptic plasticity at Golgi cell synapses. Here we have used whole-cell patch-clamp recordings and calcium imaging to investigate long-term synaptic plasticity at excitatory synapses impinging on Golgi cells. In acute mouse cerebellar slices, mossy fiber theta-burst stimulation (TBS) could induce either long-term potentiation (LTP) or long-term depression (LTD) at mossy fiber-Golgi cell and granule cell-Golgi cell synapses. This synaptic plasticity showed a peculiar voltage-dependence, with LTD or LTP being favored when TBS induction occurred at depolarized or hyperpolarized potentials, respectively. LTP required, in addition to NMDA channels, activation of T-type Ca^2+^ channels, while LTD required uniquely activation of L-type Ca^2+^ channels. Notably, the voltage-dependence of plasticity at the mossy fiber-Golgi cell synapses was inverted with respect to pure NMDA receptor-dependent plasticity at the neighboring mossy fiber-granule cell synapse, implying that the mossy fiber presynaptic terminal can activate different induction mechanisms depending on the target cell. In aggregate, this result shows that Golgi cells show cell-specific forms of long-term plasticity at their excitatory synapses, that could play a crucial role in sculpting the response patterns of the cerebellar granular layer.

**Significance statement:** This paper shows for the first time a novel form of Ca^2+^ channel-dependent synaptic plasticity at the excitatory synapses impinging on cerebellar Golgi cells. This plasticity is *bidirectional* and *inverted* with respect to NMDA receptor-dependent paradigms, with LTD and LTP being favored at depolarized and hyperpolarized potentials, respectively. Furthermore, LTP and LTD induction requires differential involvement of *T-ype and L-type voltage-gated Ca*^*2*+^ *channels* rather than the NMDA receptors alone. These results, along with recent computational predictions, support the idea that Golgi cell plasticity could play a crucial role in controlling information flow through the granular layer along with cerebellar learning and memory.

## Introduction

Several forms of plasticity are thought to provide the substrate for learning and memory in brain microcircuits (Bliss et al., 2014; Volianskis et al., 2015). In the cerebellum, which is involved in motor learning and cognitive processing (Marr, 1969; Ito, 2008; Koziol et al., 2014; Sokolov et al., 2017; D’Angelo, 2019), more than 15 diverse forms of long-term synaptic and non-synaptic plasticity have been reported at several sites across the granular layer, molecular layer and deep cerebellar nuclei (Hansel et al., 2001; Gao et al., 2012; D’Angelo et al., 2016). In the granular layer, non-synaptic plasticity has been shown both in granule cells (Armano et al., 2000) and Golgi cells (Hull et al., 2013), while long-term synaptic plasticity has been reported at the synapses made by mossy fibers with granule cells (Armano et al., 2000; Medina and Mauk, 2000; Zhang and Linden, 2006; D’Errico et al., 2009; Pugh and Raman, 2009; D’Angelo, 2014; Sgritta et al., 2017; Moscato et al., 2019). However, long-term synaptic plasticity elicited by mossy fiber stimulation in Golgi cells has not been investigated yet.

Golgi cells are the main inhibitory interneurons of the granular layer, where they receive excitatory inputs from mossy fibers and granule cells (Palay, 1974). The mossy fibers make synapses on Golgi cell basolateral dendrites inside the cerebellar glomeruli, which also contact the dendrites of granule cells. The granule cells, in turn, make synapses both on the Golgi cell basolateral dendrites through their ascending axons and on the apical dendrites through the parallel fibers. All these excitatory synapses are glutamatergic and express AMPA and NMDA receptors activated during synaptic transmission (Misra et al., 2000; Cesana et al., 2013). Golgi cells also receive inhibitory innervation from neighboring Golgi cells and other inhibitory interneurons (Dieudonne, 1998; Bureau et al., 2000; Misra et al., 2000; Hull and Regehr, 2012). Finally, Golgi cell axons contact granule cell dendrites inside the glomeruli, thus regulating information flow to the cerebellar cortex trough a mix of feedback and feed forward inhibition (D’Angelo, 2008; Kanichay and Silver, 2008; Cesana et al., 2013; D’Angelo et al., 2013). It has been suggested that Golgi cells dynamically control the gain and the temporal pattern of granule cell discharge in response to mossy fiber activity (Marr, 1969; Mitchell and Silver, 2003; D’Angelo and De Zeeuw, 2009; Billings et al., 2014) and this essential role has been supported by their acute ablation, which causes severe motor deficits and ataxia (Watanabe et al., 1998).

Golgi cells are low-frequency pacemakers (Dieudonne, 1998; Forti et al., 2006) and they have recently been shown to express Ca^2+^ channels in the dendrites (Rudolph et al., 2015). The Ca^2+^ channels, along with postsynaptic NMDA receptors, may actually enable the induction of long-term synaptic plasticity, as observed at other synapses (Lisman, 1989; Shouval et al., 2002; Volianskis et al., 2015; Leresche and Lambert, 2017). In this work we have combined patch-clamp recordings and calcium imaging techniques to show, for the first time, the existence of bidirectional long-term plasticity at the excitatory Golgi cell synapses activated by patterned mossy fiber stimulation. These results support recent computational predictions about the critical role that such forms of plasticity would play in controlling learning and computation in the cerebellar granular layer (Schweighofer et al., 2001; Garrido et al., 2013; Garrido et al., 2016).

## Methods

The experiments have been performed on 16-to 21-day-old (P0=day of birth) GlyT2-GFP mice (of either sex) heterozygous for the bacterial artificial chromosome insertion of EGFP under the glycine transporter type 2 gene (Zeilhofer et al., 2005). All procedures were conducted in accordance with European guidelines for the care and use of laboratory animals (Council Directive 2010/63/EU), and approved by the ethical committee of Italian Ministry of Health (628/2017-PR).

The mice were anesthetized with halothane (Sigma-Aldrich) and killed by decapitation in order to remove the cerebellum for acute slice preparation according to established techniques (Forti et al., 2006; Cesana et al., 2013).

### Slices preparation and solutions

The cerebellar vermis was isolated and fixed on the vibroslicer’s stage (Leica VT1200S; Leica Biosystems) with cyano-acrylic glue. Acute 220 μm-thick slices were cut in the parasagittal plane and immersed in ice-cold (2–3°C) solution containing (in mM): potassium gluconate 130, KCl 15, ethylene glycol-bis (β-aminoethyl ether) N,N,N’,N’-tetraacetic acid (EGTA) 0.2, N-2-hydroxyethyl piperazine-N-2-ethanesulphonic acid (Hepes) 20, glucose 10, pH 7.4 with NaOH (Dugué et al., 2005). Slices were incubated for at least 1 h before recordings in oxygenated bicarbonate-buffered (Kreb’s solution) saline maintained at 32°C, containing (in mM): NaCl 120, KCl 2, MgSO_4_ 1.2, NaHCO_3_ 26, KH_2_PO_4_ 1.2, CaCl_2_ 2, glucose 11 (pH 7.4 when equilibrated with 95% O_2_–5% CO_2_). During recordings, slices were placed in a chamber continuously perfused at a rate of 1.5 ml/min with oxygenated Kreb’s solution and maintained at 32°C with a Peltier feedback device (TC-324B, Warner Instrument Corp.). SR 95531 (gabazine) and strychnine were routinely added to the bath solution to block GABAergic and glycinergic inhibition, respectively.

### Electrophysiological recordings

Slices were visualized under an upright epifluorescence microscope (Axioskop 2 FS; Carl Zeiss) equipped with a 63, 0.9 NA water-immersion objective (Olympus). Whole-cell patch-clamp was performed from the soma of Golgi cells Patch pipettes were pulled from borosilicate glass capillaries (Sutter Instruments) and filled with an intracellular solution containing (in mM): potassium gluconate 145, KCl 5, HEPES 10, EGTA 0.2, MgCl_2_ 4.6, ATP-Na_2_ 4, GTP-Na_2_ 0.4, adjusted at pH 7.3 with KOH. In a different series of recordings, EGTA was increased to 10 mM. Pipettes had a resistance of 3–5 MΩ when immersed in the bath. For experiments combining current-clamp and fluorescence Ca^2+^ imaging, the pipette solution was the following (in mM): potassium gluconate 145, KCl 5, HEPES 10, MgCl_2_ 4.6, ATP-Na_2_ 4, GTP-Na_2_ 0.4 and 0.2 Oregon Green BAPTA-1 (OG1), pH adjusted at 7.3 with KOH. In a set of experiments, EGTA (0.2 mM) was substituted with BAPTA (0.2 mM).

Cell current and voltage were recorded with Multiclamp 700B, sampled with Digidata 1550 interface, and analyzed off-line with pClamp10 software (Molecular Devices). In voltage-clamp, the recorded currents were low-pass filtered at fc =10 kHz (−3dB) and digitized at 50 kHz. According to (Forti et al., 2006) recordings were discarded when the basal current at −70 mV was negative to - 150 pA. Series resistance (R_s_) was 8.3 ± 0.5 MΩ, (n=99), was constantly monitored during recordings and compensated by 40–80%. Recordings were accepted when Rs showed variations <= ±20%.

To elicit Golgi cell EPSCs the mossy fiber bundle was stimulated using a large-bore patch pipette connected to a stimulus isolation unit and filled with Kreb’s solution. Individual stimuli were 200 μs monopolar square pulses. The stimulation pipette was positioned in the white matter in an appropriate position to evoke monosynaptic mossy fiber inputs along with dysynaptic granule cell inputs according to (Cesana et al., 2013) (Fig.1A).

**Figure 1.**
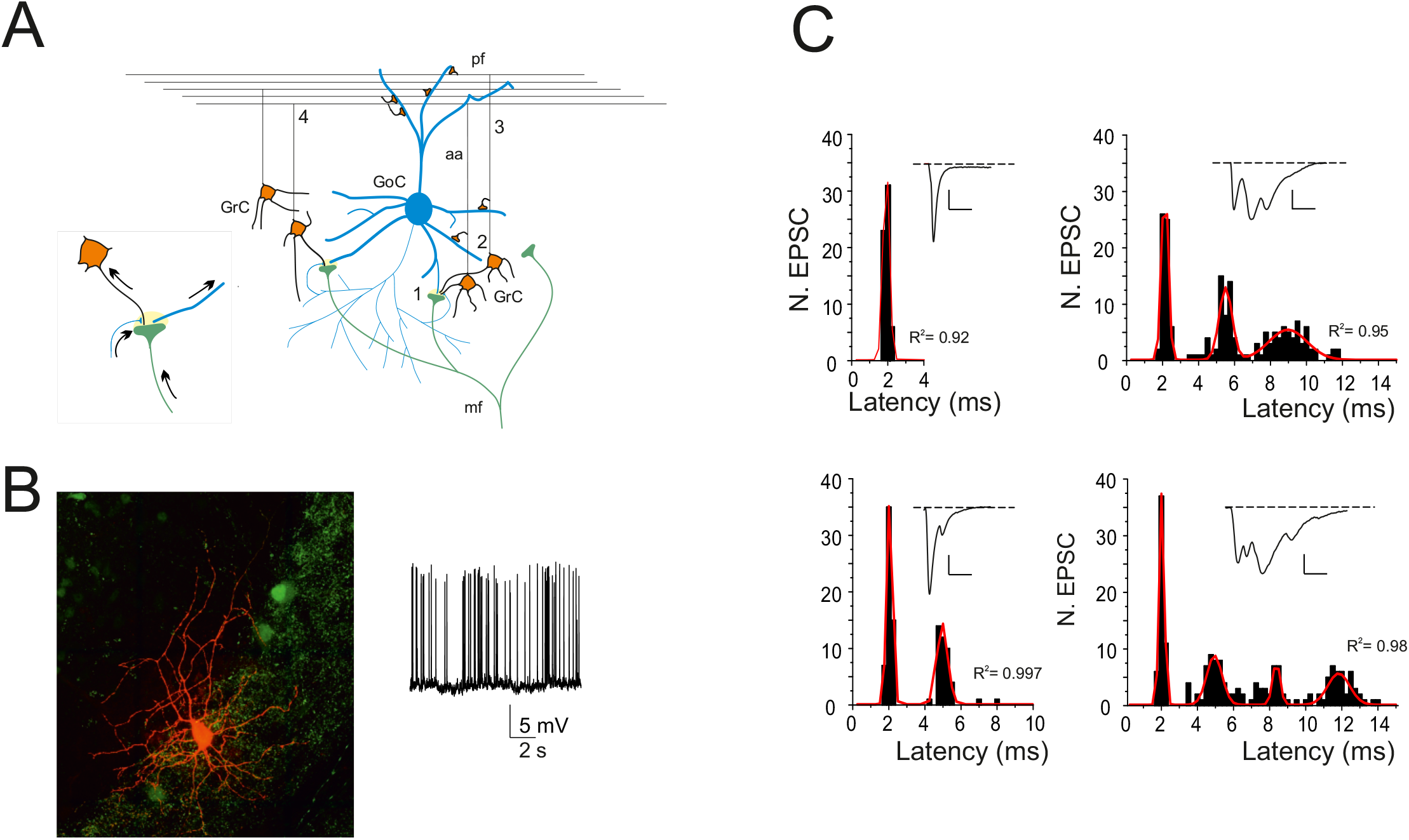
Golgi cells and their excitatory input. (A) Schematic of the afferent excitatory connections to a Golgi cell activated by mossy fiber stimulation. Golgi cell (GoC); mossy fiber (mf); granule cell (GrC); ascending axon (aa); parallel fiber (pf). The figure highlights the interactions of elements in the cerebellar glomerulus and the location of afferent Golgi cell synapses. (1) mf-Golgi cell monosynaptic connection; (2) mf-GrC-GoC disynaptic connection on aa; (3,4) short and long mf-GrC-GoC disynaptic connection on pf. The inset shows an enlargement of the contacts within the cerebellar glomerulus. (B) Fluorescence image of a Golgi cell filled with AlexaFluo-594 and recorded in whole-cell patch-clamp configuration (courtesy of Prof. Javier De Felipe). The trace on the right shows the typical Golgi cells autorhythmic firing (4.0 Hz) in the presence of GABAA and glycine receptor blockers. (C) EPSC latency in four different Golgi cells. Latency histograms show a multimodal distribution, with a first narrow peak corresponding to monosynaptic activation that could be followed by 2-3 well distinguishable later peaks corresponding to disynaptic activation (cf. panel A). A multi Gaussian fitting (red traces) is superimposed and the nonlinear regression coefficient (R^2^) are is indicated. Inset traces show the average of 30 EPSCs (scale bar 40 pA, 5 ms).

### Post hoc visualization of AlexaFluo594-filled Golgi cells

After recordings, cerebellar slices were fixed at 4°C for at least 24h in phosphate buffer saline (PBS) containing 4% paraformaldeyde (PAF), thoroughly washed in PBS and mounted in Fluoroshield Mounting Medium with DAPI (Abcam). Images were captured using a LEICA TCS SP5 confocal microscope (Centro Grandi Strumenti, UNIPV). Z-stack images were acquired for Golgi cell visualization and morphological reconstruction with the Neurolucida System (MBF Bioscience).

### Calcium imaging

Calcium imaging was performed as reported previously (Gall et al., 2005; D’Errico et al., 2009; Sgritta et al., 2017) by using Oregon green BAPTA-1 (OG1, Molecular Probes). Briefly, 200 μM OG1 was added to the intracellular solution as a substitution for the EGTA buffer. Golgi cells were identified with a ×63, 0.9 NA water immersion objective. Digital fluorescence images were obtained using an excitation light source from T.I.L.L. Photonics (Planegg) controlled through Axon Imaging Workbench AIW5.2 (INDEC Systems). Images were acquired with a 50 ms exposure/image at video rate. Acquisition started after allowing > 2 min for dye loading in the neuron. After this time, the resting fluorescence (F_0_) varied by less than 5% in each analyzed cell region for the entire recording time and the background fluorescence (B_0_) was also stationary. All stimulation protocols were separated by a minimum of 60 s in order to allow [Ca^2+^]_i_ to return to basal level. Cell damage was identified by the following signs: the failure of 200 ms depolarization at 0 mV in voltage clamp to elicit a fluorescence transient or the sudden inability of fluorescence levels to recover to baseline after stimulation. We never observed bleaching of OG1 basal fluorescence during individual stimulations. Stimulus-induced fluorescence changes were analyzed off-line in the regions of interest (ROIs). For each experiment, regions were drawn by eye defining the ROIs in the first image of a sequence, thus giving a set of two-dimensional arrays of pixels. In addition, background fluorescence was evaluated by defining a background area of similar size close to the cell. For each ROI, a measurement of the relative change in fluorescence during cell stimulation, ΔF/F_0_ (F_0_ is the mean resting fluorescence), was obtained as follows. (a) For each consecutive n^th^ image in the sequence, the fluorescence intensity f_(n)_ was evaluated in the ROI. (b) Background fluorescence was measured simultaneously in the background area, B_(n)_. Care was taken to check that background fluorescence was stationary. (c) The background-subtracted fluorescence F_(n)_ = f_(n)_ – B_(n)_ was then used to evaluate ΔF/F_0(n)_ = (F_(n)_ - F_0_)/F_0_, where F_0_ is the average background-subtracted resting fluorescence over four consecutive images before applying the stimulus. This background subtraction procedure was used to account for slice autofluorescence and/or fluorescence arising from outflow of dye from the pipette prior to seal formation. ROIs for analysis of somatic signals were chosen near the visible soma border to minimize the unfavourable surface/volume ratio for estimation of near-membrane Ca^2+^ changes. Analysis of images was performed with AIW-5.2 software.

### Drug application

Strychnine hydrochloride (1 μM) was obtained from Sigma-Aldrich. All other drugs were from Abcam: D-2-amino-5-phosphonovalerate (D-APV, 50 μM), (S)-α-Methyl-4-carboxyphenylglycine (S-MCPG, 500 μM), mibefradil dihydrochloride (10 μM), nifedipine (20 μM) and 6-imino-3-(4-methoxyphenyl)-1(6H)-pyridazinebutanoic acid hydrobromide (SR95531, gabazine, 10 μM). Stock solutions were prepared in water and stored at −20°C. During experiments, aliquots were diluted in Kreb’s solution and bath-applied.

### EPSC and TBS analysis

Post-synaptic currents (EPSCs) elicited at 0.1 Hz (test frequency) were averaged and digitally filtered at 1.5 kHz off-line. EPSC amplitude was measured as the difference between EPSC peak and baseline. After evoking EPSCs at −70 mV at the test frequency for 10-15 min (control period), the recording was switched to current clamp. Synaptic plasticity was induced by a theta-burst stimulation (TBS; eight 100 ms, 100 Hz bursts of impulses repeated every 250 ms) from an average membrane potential (V_hold_) around either −55 mV or −40 mV for LTP and LTD induction, respectively (Table 1).

**Table 1.** Membrane potential before TBS (V_hold_) between different conditions

| | LTP<br>$V_{\text{hold}}$ (mV) | LTD<br>$V_{\text{hold}}$ (mV) |
| --- | --- | --- |
| Control | -57.2 ± 1.1 (n=6) | -41.2 ± 3.4 (n=7) |
| EGTA | -59.3 ± 0.8 (n=5) | -43.2 ± 1.0 (n=5) |
| APV | -54.6 ± 1.4 (n=5) | -40.5 ± 2.0 (n=7) |
| BAPTA | -59.1 ± 3.0 (n=6) | -45.9 ± 1.5 (n=6) |
| Ca<sup>2+</sup>-imaging | -60.7 ± 1.6 (n=8) | -45.8 ± 1.1 (n=7) |
| Nifedipine | | -43.5 ± 1.1 (n=5) |
| Mibefradil | -58.2 ± 0.9 (n=6) | |

LTP and LTD $V_{\text{hold}}$ were not significantly different between different conditions.
One-way ANOVA: $F_{\text{LTP}}(5,31) = 1.58$ , $p = 0.19$ ; $F_{\text{LTD}}(5,31) = 1.55$ , $p = 0.2$

After delivering the TBS, voltage-clamp at −70 mV was reestablished and stimulation was restarted at the test frequency. The efficiency of Golgi cell synaptic excitation during TBS was expressed as mean burst depolarization (Vmean) and spike frequency (fTBS). V_mean_ was estimated as the mean of average values measured in the 100 msec of each burst (tracings were filtered at 100 Hz). V_jump_ was measured as the difference between V_hold_ and V_mean_ (see Fig.2). Long-term synaptic changes were measured after 30 min. In order to study the latency distributions of evoked EPSCs (see Fig. 1), synaptic currents were detected in the 15 ms window after a stimulus using a threshold-above-baseline detector (MiniAnalysis program; Synaptosoft Inc, Fort Lee, NJ), and their latency from stimulus onset was measured. Fittings for the latency histograms were made by using nonlinear curve fit (Gauss) (OriginPro 8). To demonstrate the presence of an evoked dysynaptic response, the probability that spontaneous EPSCs (sEPSCs) occured by chance in the time window between 2.5 and 15 ms after the stimulus (see Fig. 1C) was calculated by Poisson distribution (Forti et al., 2006)

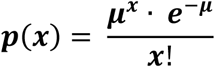

where *μ* is the average number of sEPSC per interval and *x* is the number of times an sEPSC occurs in the same interval, where *μ* is f_sEPSC_·0.0125sec [the frequency of sEPSCS is f_sEPSC_ =2.98 ± 0.25 Hz, n = 99)] In histograms shown in Fig.1, the probability of having one sEPSCs per trial was therefore *p(1)*=0.036, *p(2)*=7×10^−4^, *p(3)*= 8×10^−6^, *p(4)*=8×10^−8^, ruling out the *de facto* the potential contribution of sEPSCs.

**Figure 2.**
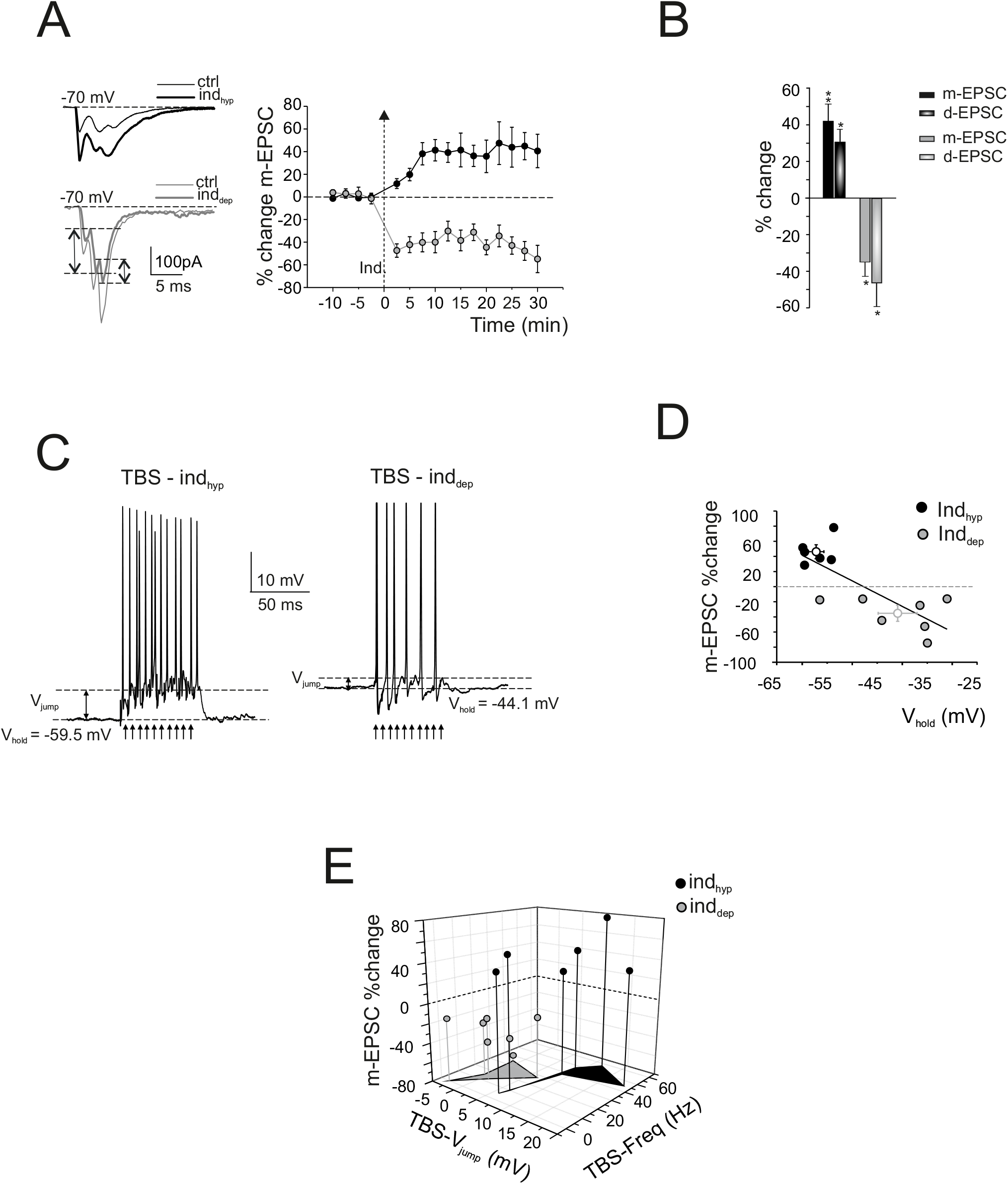
Bidirectional plasticity at Golgi cell excitatory synapses: dependence on membrane potential. LTP or LTD were induced by delivering TBS from different Golgi cell membrane potentials. (A) Left, EPSCs (average of 30 tracings) recorded before and 30 min after the delivery of TBS_hyp_ (black traces) and TBS_dep_ (gray traces). Right, average time course of m-EPSC amplitude changes during LTP and LTD. The arrow indicates the induction time and each point is the average of 15 contiguous EPSC amplitudes. Data are reported as mean ± SEM. (B) The histogram shows the average m-EPSC and d-EPSC % change following TBS_hyp_ (black) and TBS_dep_ (gray). Data are reported as mean ± SEM, *p<0.05, **p<0.01, Student’s paired *t* test. (C) Golgi cell response during TBS_hyp_ and TBS_dep_. Note the stronger depolarization and spike generation in TBS_hyp_ than TBS_dep_. The figure shows V_hold_ and V_jump_ in the two cases. (D) The plot shows the relationship between membrane holding potential (V_hold_) and m-EPSC amplitude and can be fitted with straight line (linear regression R^2^ =0.57, n=13, p(F)=0.002) (E) The 3D graph shows m-EPSC amplitude change as a function of the efficiency of Golgi cell synaptic excitation during TBS_hyp_ and TBS_dep_ evaluated as V_jump_ and spike frequency.

### Statistical procedures

All data are reported as mean ± MSE. The normality of data was checked by applying the Shapiro-Wilk’s test. Means were compared by a Student’s *t*-test or by one-way parametric analysis of variance (ANOVA). Where appropriate, data were further assessed by conducting the Tukey *post hoc* test. The analysis was two-sided, with level of significance α = 0.05. All statistical analyses were done using OriginPro 8.

## Results

Whole-cell patch-clamp recordings were performed from Golgi cells in acute cerebellar slices of GlyT2-GFP mice (P16-P21). Golgi cells were identified as GFP fluorescent neurons in the granular layer showing a large soma (>10 μm diameter) and typical single-spike rhythmic firing (1-10 Hz; Fig.1B) (Forti et al., 2006; Cesana et al., 2013) from an average membrane potential V_m_ = - 49.8 ± 0.8 mV (n=99). It should also be considered that, in addition to Golgi cells, GFP is expressed in other types of GABAergic interneurons of the granular layer of GlyT2-GFP mice, namely Lugaro cells and globular cells (Zeilhofer et al., 2005; Simat et al., 2007), which cannot be easily identified in acute slices due to the high density and extensive overlap among the numerous GFP-containing axonal and dendritic processes. In a subset of cells (n=8), we obtained *post hoc* morphological confirmation by filling interneurons with 50 μM AlexaFluo 594 and carrying out laser confocal microscopy reconstructions (Fig.1B). Golgi cells showed a soma diameter of 17±0.833 μm (n=8) with several basolateral dendrites remaining in the granular layer (30.85±6.15 μm long; n= 8) and apical dendrites (70.14±9.73 μm long; n=8) ascending towards the molecular layer. Their axons diffusely extended in the granular layer (Dieudonne, 1998; Misra et al., 2000).

### EPSCs evoked in Golgi cells by mossy fiber stimulation

In order to investigate the contributions of mossy fiber and granule cell synaptic inputs to Golgi cell excitation, EPSCs were recorded at −70 mV in the presence of gabazine (10 μM) and strychnine (1 μM) to block GABAergic and glycinergic synapses, respectively (Dumoulin et al., 2001; Hull and Regehr, 2012; Cesana et al., 2013). Golgi cells EPSCs were activated by mossy fiber stimulation at least 300 μm away from soma to avoid direct activation of granule cell axons (Cesana et al., 2013). This configuration recruited monosynaptic mossy fiber inputs and disynaptic mossy fiber - granule cell inputs (Fig. 1A,C). Mossy fiber stimulation, adjusted to obtain 50-250 pA responses, evoked in a group of Golgi cells (18 out of 99) an isolated EPSC with short latency (Fig. 1C), with latency histograms showing a single narrow peak at 2.10 ± 0.13 ms (n=18). In the remaining cells (81 out of 99), stimulation evoked multiple EPSCs and gave rise to multiple components in the average trace (Fig. 1C). A Poisson distribution generated with the mean sEPSC frequency ruled out that these late events were actually sEPSCs (see Methods). The latency histograms of multimodal distributions showed a first narrow peak at 2.05 ± 0.05 ms (n=81), followed by late humps [peaking at 4.64 ± 0.19 ms (n=33), 6.65 ± 0.27 ms (n=40), 10.45 ± 0.89 (n=7), respectively]. The multiple peaks were separated by 2.30 ± 0.13 ms (n=67), 2.65 ± 0.29 ms (n=33) and 3.55 ± 0.76 ms (n=7), respectively. In aggregate, synaptic responses could be separated into monosynaptic EPSCs (m-EPSCs) and disynaptic EPSCs (d-EPSCs). According to a previous detailed analysis (Cesana et al., 2013), the m-EPSCs are likely to arise at the mossy fiber – Golgi cell synapse (Fig. 1A, connection 1), whereas d-EPSC are likely to originate from mossy fiber - granule cell - Golgi cell disynaptic inputs on the ascending axon (Fig. 1B, connection 2) as well as from short and long mossy fiber - FGFgranule cell - Golgi cell disynaptic inputs on the parallel fibers (Fig. 1B, connection 3 and 4).

### Voltage-dependent plasticity at mf-Golgi cell synapses

The presence of long-term synaptic plasticity between mossy fibers and Golgi cells was investigated in the presence of GABA_A_ and glycine receptor blockade by 10 μM gabazine and 1 μM strychnine, respectively (Fig.2). Following a 10-min control period to monitor recording stability, mossy fibers were stimulated with high-frequency impulse trains in current-clamp reproducing a TBS patterns (See Methods), which is known to induce persistent neurotransmission changes at the neighboring synapses made by mossy fibers with granule cells (D’Angelo et al., 1999; Hansel et al., 2001; D’Errico et al., 2009; Andreescu et al., 2011). At the mossy fiber - granule cell synapse, membrane potential before TBS (V_hold_) is an important determinant of the sign of synaptic plasticity, with LTP arising from depolarized and LTD from hyperpolarized potentials. Here too, TBS was delivered from different levels of V_hold_ (depolarized and hyperpolarized, TBS_dep_ and TBS_hyp_).

Quite unexpectedly, following TBS pairing at V_hold_ = −40.9 ± 3.4 mV (n=7), both m-EPSC and d-EPSC amplitude decreased by −35.2 ± 8.6 % (n=7; Student’s paired *t* test, p=0.036) and −47.1 ± 12.82 % (n=6; Student’s paired *t* test, p=0.048). Conversely, following TBS pairing at V_hold_ = - 57.2 ± 1.2 mV (n=6; this potential was significantly different from the other one; Student’s unpaired *t* test, p=0.002), m-EPSC and d-EPSC amplitude increased by 46.2 ± 7.2% (n=6; Student’s paired *t* test, p=0.007) and 30.6 ± 8.2% (n=6; Student’s paired *t* test, p=0.049; Fig.2A,B). Therefore, the relationship between membrane potential and plasticity was inverted in Golgi cells compared to granule cells (D’Angelo et al., 1999; D’Errico et al., 2009).

The m-EPSC changes obtained at different V_hold_ showed a significant negative linear correlation (R^2^=0.57; one-way ANOVA, p=0.0018) with LTD and LTP occurring at the depolarized and hyperpolarized V_hold_, respectively (Fig. 2D). The m-EPSC changes also depended on the membrane depolarizing jump (V_jump_) during TBS (Fig. 2C,E).

In recordings that showed LTP, TBS caused a strong Golgi cell excitation (V_jump_ = 10.0 ± 1.9 mV, n=6) characterized by a depolarizing hump sustaining a robust action potential discharge (f_TBS_= 33.7 ± 8.7 Hz, n=6). Conversely, in the recordings that showed LTD, TBS was unable to generate a clear depolarizing hump (V_jump_ = −0.99 ± 0.54 mV, n=7; Student’s unpaired *t* test, p=0.0016) but could still evoke an action potential discharge (f_TBS_ = 20.0 ± 4.7 Hz, n=7). Although the average action potential discharge frequency was higher in Golgi cells showing LTP than in those showing LTD, the difference was not statistically significant (Student’s unpaired *t* test, p=0.2).

These results show an *inverted* voltage-dependent bidirectional plasticity at the mossy fiber - Golgi cell synapses.

### LTP and LTD require an elevation of postsynaptic Ca^2+^

In a different series of recordings, in order to determine whether intracellular Ca^2+^ was required for LTP and LTD at mossy fiber-Golgi cell synapses, the pipette intracellular solution was supplemented with the calcium buffer, 10 mM EGTA. After establishing the whole-cell configuration, we waited about 15 min to ensure adequate perfusion of EGTA into the dendritic compartment before applying LTP and LTD protocols. Figure 3 shows the time course of m-EPSC changes. The 10 mM EGTA prevented both LTP (m-EPSC, −4.2± 4.4%, n=5; Student’s paired *t* test, p=0.53; d-EPSC, −7.5 ± 6.0%, n=7; Student’s paired *t* test, p=0.35) and LTD (m-EPSC, 0.78± 6.8%, n=5; Student’s paired *t* test, p=0.8; d-EPSC, 6.7 ± 15.2%, n=5; Student’s paired *t* test, p=0.3) (Fig. 3A, B). It should be noted that, compared to control, TBS_hyp_ (−59.3 ± 0.8 mV, n=5; Table 1) showed a weaker Golgi cell excitation (V_jump_ = 4.4 ± 1.1 mV, n=5; Student’s unpaired *t* test, p=0.035) accompanied by reduced action potential discharge (f_TBS_ = 7.5 ± 6.3 Hz, n=5; Student’s unpaired *t* test, p=0.038; Fig. 3C), while TBS_dep_ (−43.2 ± 1.0 mV, n=5; Table 1) showed a stronger Golgi cell excitation (V_jump_ = 3.8 ± 1.6 mV, n=5; Student’s unpaired *t* test, p=0.034) accompanied by a robust action potential discharge (f_TBS_ = 39.0 ± 3.4 Hz, n=5; Student’s unpaired *t* test, p=0.008; Fig. 3C).

**Figure 3.**
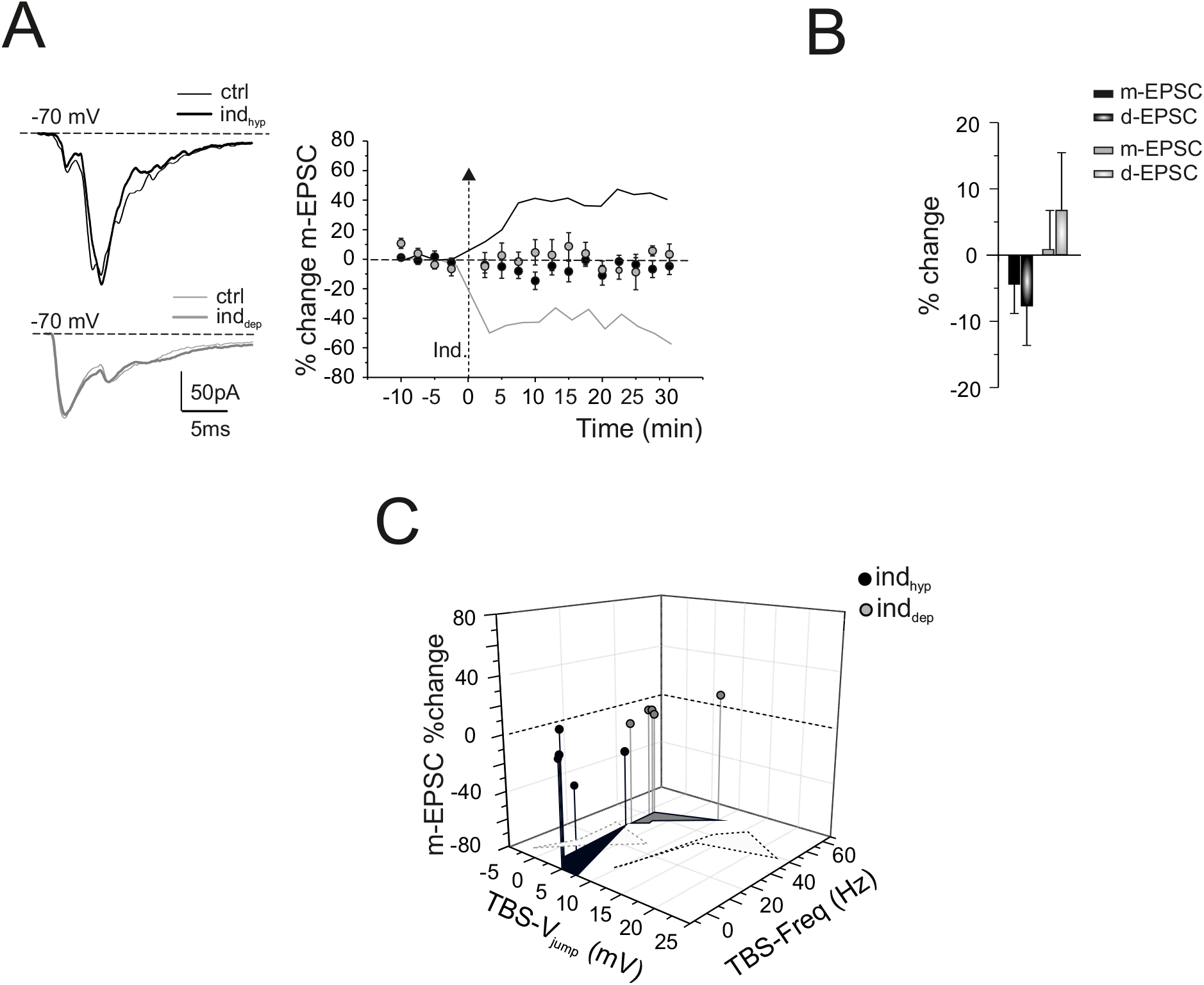
Postsynaptic Ca^2+^-dependent induction of LTP and LTD. (A) Left, EPSCs (average of 30 tracings) recorded before and 30 min after the delivery of TBS_hyp_ (black traces) and TBS_dep_ (gray traces) with 10 mM EGTA in the intracellular solution. Right, average time course of m-EPSC amplitude changes. The arrow indicates the induction time and each point is the average of 15 contiguous EPSC amplitudes. Note that high EGTA prevented both LTP and LTD. The black and gray lines are replotted from Fig. 2A to compare LTP and LTD in control condition. Data are reported as mean ± SEM. (B) The histogram shows the average m-EPSC and d-EPSC % change following TBS_hyp_ (black) and TBS_dep_ (gray). Data are reported as mean ± SEM. (C) The 3D graph shows m-EPSC amplitude change as a function of the efficiency of Golgi cell synaptic excitation during TBS_hyp_ and TBS_dep_ (evaluated both as V_jump_ and spike frequency) in the presence of high EGTA. The black and gray dotted areas are replotted from Fig. 2E to compare m-EPSC amplitude change as a function of the efficiency of Golgi cell synaptic excitation during TBS_hyp_ and TBS_dep_ in control.

These results indicate that long-term plasticity at Golgi cell excitatory synapses requires postsynaptic calcium changes for induction, like other forms of LTP and LTD at central synapses (Artola et al., 1990; Artola and Singer, 1993; D’Angelo et al., 1999; Malenka and Bear, 2004; D’Errico et al., 2009; Sgritta et al., 2017).

### Voltage dependence of intracellular Ca^2+^ changes during induction

The relationship between [Ca^2+^]_i_ changes and Golgi cell synaptic plasticity was investigated by Ca^2+^ imaging measurements using 200 μM OG1 in the patch pipette (Gall et al., 2005; D’Errico et al., 2009; Sgritta et al., 2017). The [Ca^2+^]_i_ increase in the Golgi cell basolateral dendritic compartment (Fig. 4A,B) was significantly higher with TBS_hyp_ than TBS_dep_ [(F/F_0_)_max_ = 0.23± 0.02 vs 0.13 ± 0.02, n= 6 (13 dendrites); Student’s paired *t* test, p = 3.7e^−6^]. Since OG1 has a Ca^2+^ affinity similar to BAPTA, we performed control recordings with BAPTA in the intracellular pipette instead of EGTA allowing, therefore, a direct comparison of imaging with patch-clamp recordings. As expected, TBS_hyp_ induced LTP (m-EPSC, Student’s unpaired *t* test, p=0.98; d-EPSC, Student’s unpaired *t* test, p=0.31) and TBS_dep_ induced LTD (m-EPSC, Student’s unpaired *t* test, p=0.45; d-EPSC, Student’s unpaired *t* test, p=0.65). These LTP and LTD were indistinguishable from those observed using 200 μM EGTA (Fig. 4C, D). It should be noted that, during TBS_hyp_ and TBS_dep_, Golgi cell excitation (TBS_hyp_, Student’s unpaired *t* test, p=0.12; TBS_dep_, Student’s unpaired *t* test, p=0.5) and action potential discharge (TBS_hyp_, Student’s unpaired *t* test, p=0.2; TBS_dep_, Student’s unpaired *t* test, p=0.54) were not significantly different when using 200 μM BAPTA instead of 200 μM EGTA (Fig. 4E).

**Figure 4.**
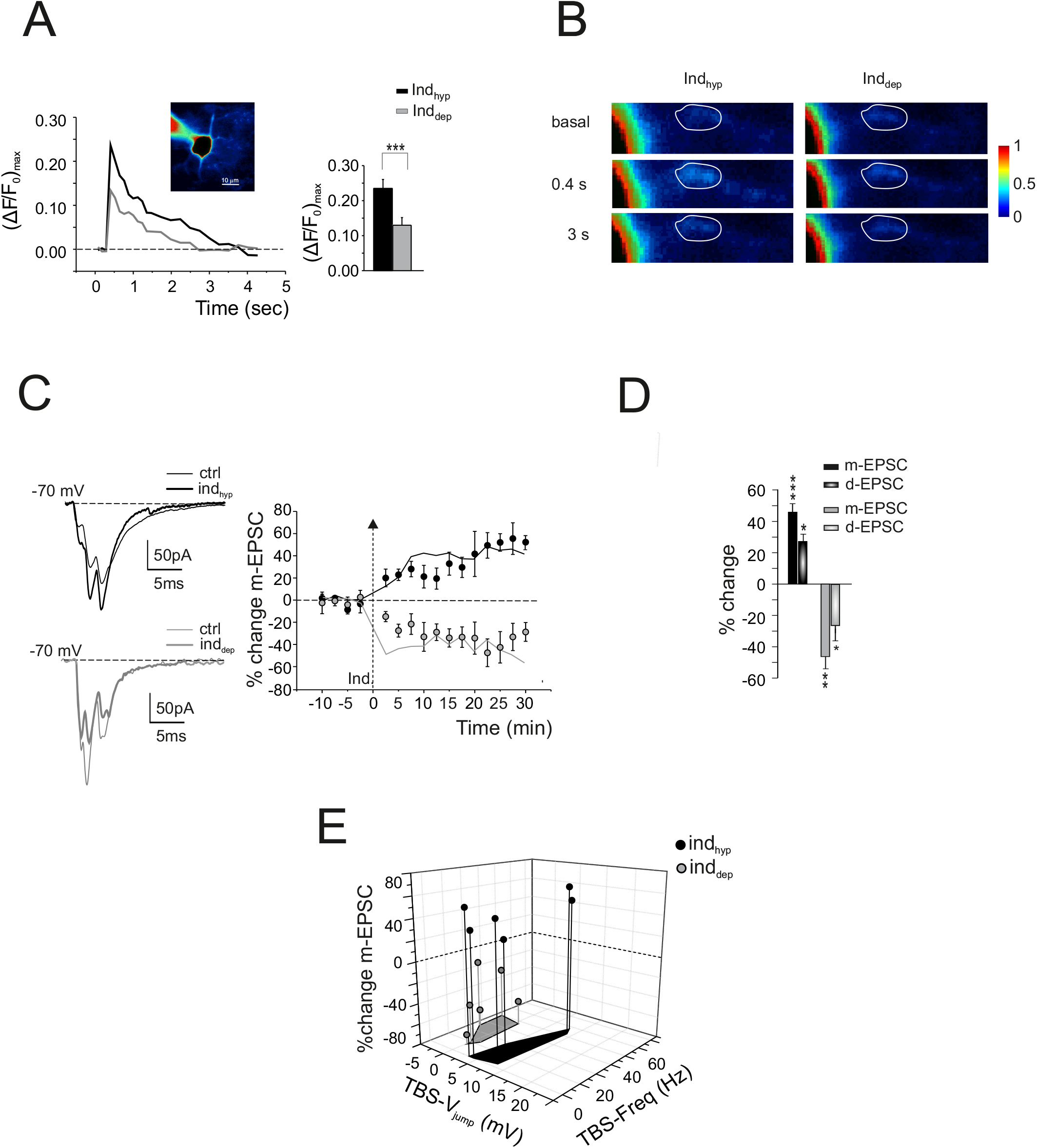
The relationship between [Ca^2+^]_i_ changes and Golgi cell synaptic plasticity. (A) Left, Ca^2+^ transients elicited by TBS_hyp_ (black traces) and TBS_dep_ (gray traces). Inset, unprocessed fluorescence image of a Golgi cell filled with 200 μM OG1. Scale bar, 10 μm. Fluorescence intensity is color coded with arbitrary units within an interval chosen to allow the visualization of the basolateral dendrites (note that, although the soma fluorescence appears saturated within this interval, it does not saturate the CCD detectors). Right, histograms show the (ΔF/F_0_)_max_ induced by TBS_hyp_ and TBS_dep_, respectively. Data are reported as mean ± SEM, ***p<0.005, Student’s paired *t* test. (B) The time series of pseudoratio images from the same Golgi cell in A (warmer color code for higher ΔF/F_0_) is shown. ΔF/F_0_ is evaluated in the ROI. (C) Left, EPSCs (average of 30 tracings) recorded before and 30 min after the delivery of TBS_hyp_ (black traces) and TBS_dep_ (gray traces) with 200 μM BAPTA in the intracellular solution. Right, the average time course of m-EPSC amplitude changes. The arrow indicates the induction time and each point is the average of 15 contiguous EPSC amplitudes. LTD and LTP observed using 200 μM BAPTA was indistinguishable from that observed in control condition (0.2 mM EGTA). The black and gray lines are replotted from Fig. 2A to compare LTP and LTD in control condition. Data are reported as mean ± SEM. (D) The histogram shows the average m-EPSC and d-EPSC % change following TBS_hyp_ and TBS_dep_. Data are reported as mean ± SEM, *p<0.05, **p<0.01, ***p<0.005, Student’s paired *t* test. (E) The 3D graph shows m-EPSC amplitude change as a function of the efficiency of Golgi cell synaptic excitation during TBS_hyp_ and TBS_dep_ (evaluated both as V_jump_ and spike frequency)

### The role of NMDA receptors in LTP and LTD induction

The classical mechanism by which membrane depolarization influences the induction of synaptic plasticity passes through the removal of voltage-dependent Mg^2+^ block from NMDA channels (Bliss et al., 2014) To examine whether NMDA receptors actually play a role in plasticity at the excitatory Golgi cell synapses, we delivered TBS in the presence of an NMDA receptor antagonist, D-APV. Bath application of D-APV (50 μM) did not produce any remarkable changes in basal transmission (m-EPSC, −3.3 ± 3.1%, n=13;, Student’s paired *t* test, p=0.16 and d-EPSC, −9.7 ± 4.9%, n=13; Student’s paired *t* test, p=0.13; Fig. 5A,B) but prevented the induction of LTP by TBS_hyp_ (V_hold_ = −54.6 ± 1.4 mV; n=6; Table 1). m-EPSC and d-EPSC amplitude changed by −8.2 ± 7.4 % (n=6; Student’s paired *t* test, p=0.18) and −3.0 ± 20.0 % (n=7; Student’s paired *t* test, p=0.7; Fig. 5C, D), respectively. TBS_hyp_ showed a weaker Golgi cell excitation compared to control (V_jump_ = 3.6 ± 0.8 mV, n=6; Student’s unpaired *t* test,p=0.018) accompanied by a reduced action potential discharge (f_TBS_ = 2.5 ± 1.6 Hz, n=6; Student’s unpaired *t* test,p=0.015; Fig. 5E). Conversely, after TBS_dep_ (V_hold_ = −40.2 ± 1.7 mV; n=7; Table 1), m-EPSC and d-EPSC amplitude were reduced to 26.1 ± 9.5 % (n=7; Student’s paired *t* test, p=0.037) and 19.4 ± 6.7 % (n=7; Student’s paired *t* test, p=0.05; Fig. 5C, D), respectively. The magnitude of LTD in the presence of D-APV was not significantly different from control (Student’s unpaired *t* test, p=0.49). Likewise, Golgi cell excitation (Student’s unpaired *t* test, p=0.7) and action potential discharge during TBS_dep_ were not different from control (Student’s unpaired *t* test, p=0.73) (Fig. 5E). These results indicate that NMDA receptor activation is required for LTP but not for LTD at Golgi cell excitatory synapses.

**Figure 5.**
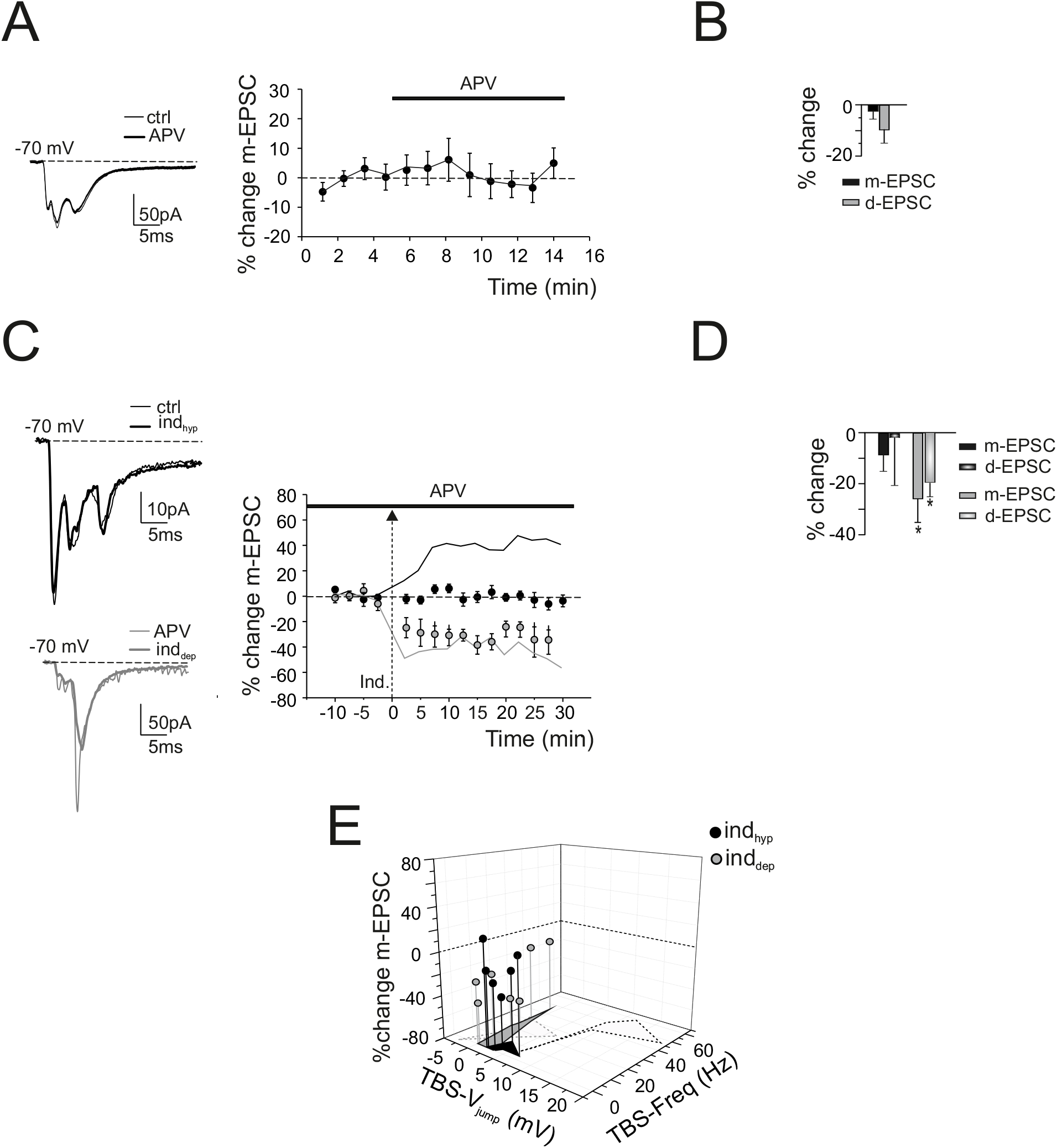
NMDA receptor-dependent induction of LTP. (A) Left, EPSCs (average of 30 tracings) recorded before and 10 min after the bath application of the NMDA receptor antagonist, D-APV (50 μM). Right, the average time course of m-EPSC amplitude changes. The black bar indicates the duration of perfusion and each point is the average of 5 contiguous EPSC amplitudes. Note that D-APV did not cause changes in basal transmission. (B) The histogram shows the average m-EPSC and d-EPSC % change in the presence of D-APV. Data are reported as mean ± SEM. (C) Left, EPSCs (average of 30 tracings) recorded before and 30 min after the delivery of TBS_hyp_ (black traces) and TBS_dep_ (gray traces) with 50 μM D-APV in the bath (black bar). Right, the average time course of m-EPSC amplitude changes. The arrow indicates the induction time and each point is the average of 15 contiguous EPSC amplitudes. Note that D-APV completely blocked the induction of LTP while the magnitude of LTD was not significantly different from that in the absence of NMDA receptor block. The black and gray lines are replotted from Fig. 2A to compare LTP and LTD in control condition. Data are reported as mean ± SEM. (D) The histogram shows the average m-EPSC and d-EPSC % change following TBS_hyp_ and TBS_dep_ in the presence of D-APV. Data are reported as mean ± SEM, *p<0.05, Student’s paired *t* test. (E) The 3D graph shows m-EPSC amplitude change as a function of the efficiency of Golgi cell synaptic excitation during TBS_hyp_ and TBS_dep_ (evaluated both as V_jump_ and spike frequency) in the presence of D-APV. The black and gray dotted areas are replotted from Fig. 2E to compare m-EPSC amplitude change as a function of the efficiency of Golgi cell synaptic excitation during TBS_hyp_ and TBS_dep_ in control condition.

### The role of voltage-gated calcium channels in LTP and LTD induction

Since Golgi cell depolarization activates voltage-gated ionic channels, and since both LTP and LTD proved to depend on postsynaptic membrane voltage and Ca^2+^ concentration changes, we examined the role of voltage-gated calcium channels in the induction of LTD and LTP at mossy fiber-Golgi cell synapses. Actually both T-type (LVA) and L-type (HVA) Ca^2+^ channels have been reported along with their subcellular distribution (Solinas et al., 2007; Rudolph et al., 2015). T-type Ca^2+^ channels activate at membrane potentials positive to −70 mV and are fully inactivated beyond - 40 mV, while L-type Ca^2+^ channels activate at membrane potentials positive to −30 mV, suggesting a specific and differential engagement in LTP and LTD.

We first evaluated the effect of the LVA Ca^2+^ channel blocker, mibefradil, on LTP. Mibefradil is a potent inhibitor of T-type Ca^2+^ (Cav3.x) currents (Mishra and Hermsmeyer, 1994; Martin et al., 2000) and their involvement in long-term synaptic plasticity has been reported (Leresche and Lambert, 2017). Bath application of 10 μM mibefradil did not cause any significant changes in basal synaptic transmission (m-EPSC, −6.5 ± 1.5%, n=6; Student’s paired *t* test, p=0.16 and d-EPSC, −15.3 ± 8.8%, n=6; Student’s paired *t* test, p=0.17; Fig. 6A,B) but prevented the induction of LTP by TBS_hyp_ (V_hold_ = −58.2 ± 0.9 mV; n=6; Table 1). m-EPSC and d-EPSC amplitude changed by −22.2 ± 9.2 % (n=6; Student’s paired *t* test, p=0.16) and −4.4 ± 10.0 % (n=6; Student’s paired *t* test, p=0.47; Fig. 6C,D), respectively. Moreover, TBS showed a weaker Golgi cell excitation compared to control (V_jump_ = 3.9 ± 0.3 mV, n=6; Student’s unpaired *t* test, p=0.024) accompanied by a reduced action potential discharge (f_TBS_ = 8.3 ± 5.06 Hz, n=6; Student’s unpaired *t* test, p=0.035; Fig. 6E).

**Figure 6.**
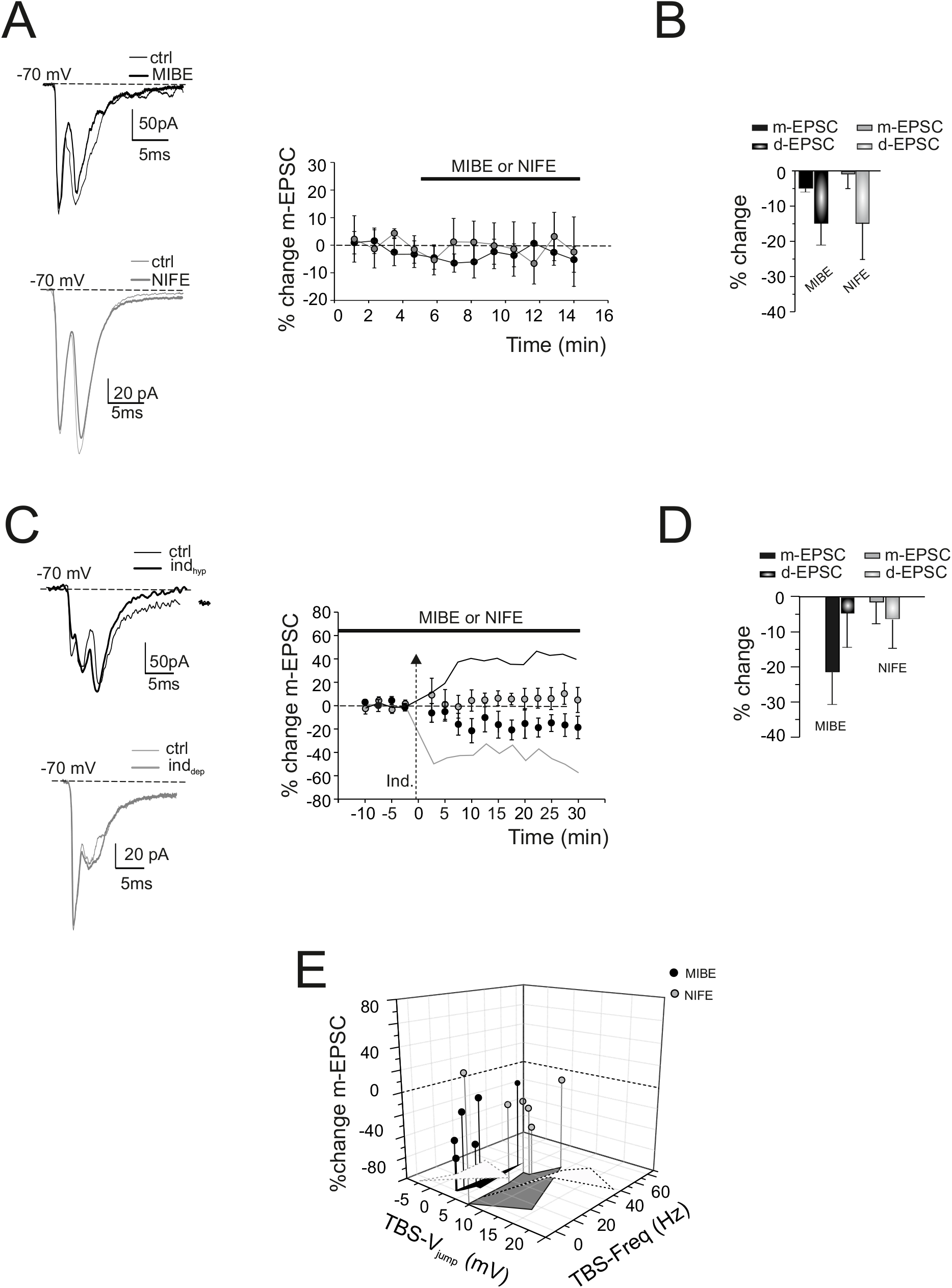
Voltage-gated calcium channel-dependent induction of LTP and LTD. (A) Left, EPSCs (average of 30 tracings) recorded before and 10 min after the bath application of the T-type and L-type Ca^2+^ channel blockers, 10 μM mibefradil and 20 μM nifedipine, respectively. Right, the average time course of m-EPSC amplitude changes. The black bar indicates the duration of perfusion and each point is the average of 5 contiguous EPSC amplitudes. Note that neither mibefradil or nifedipine produced any remarkable changes in basal transmission. (B) The histogram shows the average m-EPSC and d-EPSC % change in the presence of mibefradil and nifedipine. Data are reported as mean ± SEM. (C) Left, EPSCs (average of 30 tracings) recorded before and 30 min after the delivery of TBSh_yp_ (black traces) in the presence of 10 μM mibefradil and TBS_dep_ (gray traces) in the presence of 20 μM nifedipine in the bath (black bar). The arrow indicates the induction time and each point is the average of 15 contiguous EPSC amplitudes. Note that mibefradil blocked the induction of LTP while nifedipine blocked the induction of LTD. The black and gray lines were replotted from Fig. 2A to compare show LTP and LTD in control condition. Data are reported as mean ± SEM. (D) The histogram shows the average m-EPSC and d-EPSC % change following TBShyp in the presence of mibefradil and TBS_dep_ in the presence of nifedipine. Data are reported as mean ± SEM. (E) The 3D graph shows m-EPSC amplitude change as a function of the efficiency of Golgi cell synaptic excitation (evaluated both as V_jump_ and spike frequency) during TBS_hyp_ in the presence of mibefradil and during TBS_dep_ in the presence of nifedipine. The black and gray dotted areas are replotted from Fig. 2E to compare m-EPSC amplitude changes as a function of the efficiency of Golgi cell synaptic excitation during TBS_hyp_ and TBS_dep_ in control condition.

We then evaluated the effect of the HVA Ca^2+^ channel blocker, nifedipine, on LTD. Nifedipine is a potent inhibitor of L-type Ca^2+^ (Cav1.x) currents (Nguemo et al., 2013; Striessnig et al., 2015) and their involvement in long-term synaptic plasticity has been reported (Leresche and Lambert, 2017). Bath application of 20 μM nifedipine did not cause any significant changes in basal synaptic transmission (m-EPSC, −0.1 ± 4.1%, n=5; Student’s paired *t* test, p=0.7 and d-EPSC, −16.8 ± 1.36%, n=5; Student’s paired *t* test, p=0.7; Fig. 6A,B) but blocked the induction of LTD by TBS_dep_ (V_hold_ =-43.5 ± 1.1 mV; n=5; Table 1). m-EPSC and d-EPSC amplitude changed by −1.2 ± 7.5 % (n=5; Student’s paired *t* test, p=0.37) and −6.5 ± 9.9 % (n=5; Student’s paired *t* test, p=0.42; Fig. 6C, D), respectively. In addition, TBS showed a stronger Golgi cell excitation compared to control (V_jump_ = 5.1 ± 2.3 mV, n=5; Student’s unpaired *t* test, p=0.05) whereas action potential discharge (f_TBS_ = 26.5 ± 5.4 Hz, n=5; Student’s unpaired *t* test, p=0.42; Fig. 6E) was not significantly different. These results indicate that T-type and L-type Ca^2+^ channel activation plays a fundamental and distinctive role in the voltage-dependent induction of long-term synaptic plasticity at excitatory Golgi cell synapses.

## Discussion

This work reports for the first time long-term synaptic plasticity at excitatory synapses impinging on cerebellar Golgi cells in mice. This plasticity is bidirectional, with LTD and LTP being favored at depolarized and hyperpolarized potentials, respectively. Interestingly, this voltage-dependence of induction is inverted not just with respect to that of many other brain excitatory synapses (Lisman, 1989; Artola et al., 1990; Artola and Singer, 1993; Malenka and Bear, 2004; Feldman, 2009; Bliss et al., 2014) but also to that occurring at the neighboring granule cell synapses, which are conjointly activated with Golgi cells by mossy fibers in the cerebellar glomerulus (D’Errico et al., 2009). Thus the mossy fiber terminal can make different plasticity depending on its target cells. Correspondingly, the mechanisms of LTP and LTD induction involved differential engagement of voltage-gated Ca^2+^ channels rather than NMDA receptors alone (Volianskis et al., 2015). LTP and LTD generated by the granule cell ascending axon and parallel fibers were indistinguishable from that generated by mossy fibers, implying that mechanisms of induction reflected specific postsynaptic properties of the Golgi cell. It should be noted that, although the secondary multiple peaks in latency histograms were most likely related to distinct granule cell inputs to the basolateral and apical dendrites, it was not possible in these experiments to further dissect these components in terms of their induction properties.

### The mechanisms of inverted bidirectional plasticity

The way Golgi cell plasticity is induced is in line with a set of observations about the membrane channel complement and the physiological activity of Golgi cells. Golgi cell dendrites are endowed with Ca^2+^ channels (Rudolph et al., 2015) and the excitatory synapses express NMDA receptors (Misra et al., 2000; Cesana et al., 2013). Accordingly, mossy fiber bursts caused synaptic calcium influx and dendritic calcium spikes in a voltage-dependent manner (see Fig. 3). It is therefore conceivable that modulation of Golgi cell membrane potential would profoundly affect calcium influx through these membrane channels thereby affecting the induction of long-term synaptic plasticity. We have actually tested the effect of T-type and L-type Ca^2+^ channels that are reported to play key roles in LTP and LTD induction (Leresche and Lambert, 2017). Neither T-type Ca^2+^ channel blockade (with mibefradil) nor L-type Ca^2+^ channel blockade (with nifedipine) significantly affected basal neurotransmission, suggesting that their action was primarily postsynaptic in our experiments. Actually, T-type Ca^2+^ channel activation (along with NMDA receptors) was required for LTP but not for LTD, while L-type channel activation was required for LTD. It should be noted that a recent work has reported a differential distribution of T-type and R-type Ca^2+^ channels in distal dendrites and of L-type, N-type and P-type channels in basolateral dendrites (Rudolph et al., 2015). However, we did not find significant differences when considering monosynaptic and dysynaptic EPSCs. Subtle distinctive effects of these different Ca^2+^ channel subtypes on LTP and LTD induction remain to be investigated.

In addition to identify a correlation of membrane mechanisms with LTP or LTD, we observed that voltage changes during TBS were stronger from hyperpolarized than depolarized membrane potentials (measured with respect to the pacemaking potential of the cells). In aggregate, this set of observations can be explained by the *following model*. From slightly hyperpolarized potentials (about −57 mV), at which T-type Ca^2+^ channels are de-inactivated, TBS would cause a strong Golgi cell excitation by virtue of T-type Ca^2+^ channel opening. This would generate a depolarizing hump (actually a calcium spike) unblocking the NMDA channels and raising intracellular calcium above LTP threshold. From slightly depolarized potentials (about −41 mV), at which T-type Ca^2+^ channels are inactivated, TBS would cause weak Golgi cell excitation since the calcium spike would be prevented along with NMDA channel unblock. Nonetheless, enough calcium would still enter through L-type Ca^2+^ channels causing LTD. It is therefore possible that the intracellular calcium-dependent machinery respects the Lisman paradigm [low calcium for LTD, high calcium for LTP (Lisman, 1989; Shouval et al., 2002)] with a voltage-dependent induction mechanism enhancing calcium influx from hyperpolarized membrane potential. Similar (but probably not identical) LTP and LTD induction mechanisms engaging combinations of NMDA and voltage-gated T-type and L-type Ca^2+^ channels have also been reported at synapses in thalamic nuclei, deep cerebellar nuclei, cerebral cortex, hippocampus and striatum (Leresche and Lambert, 2017).

### Functional implications

Since the depolarization and hyperpolarization used to induced plasticity were around the pacemaking potential of the Golgi cell (about ±10 mV), it is possible that the induction mechanism is physiologically regulated by local network activity. The membrane potential change needed to switch between LTP and LTD can be driven by inhibitory and excitatory synaptic activity. While excitatory synapses are activated by mossy fibers as well as in granule cells through their ascending axons and parallel fibers (Cesana et al., 2013), inhibitory synapses are provided by neighboring Golgi cells (Hull and Regehr, 2012) and from Lugaro cells, while inputs from stellate and basket cells are still disputed (Dieudonne, 1998; Bureau et al., 2000; Misra et al., 2000). Moreover, neuromodulators could also bias Golgi cell membrane potential (Geurts et al., 2002; Schweighofer et al., 2004; Fleming and Hull, 2019). The activity of Golgi cells can also be influenced by the climbing fibers (Xu and Edgley, 2008) although the mechanisms is unclear and direct synaptic contacts have not been demonstrated (Galliano et al., 2013). There is therefore a rich set of mechanisms that could determine the direction of plastic changes at Golgi cell excitatory synapses.

This novel form of plasticity places the Golgi cells in a pivotal position to regulate information flow through the granular layer. Actually, strong mossy fiber bursts turn out to induce both LTP of glutamatergic excitation in granule cells (D’Errico et al., 2009), LTD of glutamatergic excitation in Golgi cells (this paper) and LTD of GABAergic inhibition between Golgi cells and granule cells (Mapelli et al., 2015). This would eventually cause a global long-term enhancement of granule cell excitation (both directly and indirectly through reduced Golgi cell-mediated inhibition) thereby enhancing transmission gain. The opposite would occur with poorly depolarizing inputs. Interestingly, hyperpolarization was also shown to induce a long-term increase in the spontaneous firing rate of cerebellar Golgi cells (Hull et al., 2013) and very strong impulse trains in the parallel fibers were reported to induce a form of LTD (Robberechts et al., 2010) suggesting that multiple long-term regulatory mechanisms coexist.

### Conclusions

Long-term changes in Golgi cell synaptic transmission determined by patterned mossy fiber synaptic inputs and could have relevant implications for granular layer processing. The granular layer is thought to operate as an adaptive filter (Dean and Porrill, 2010) and modifying the inhibitory effect of Golgi cells through plasticity has been proposed as a key factor to regulate signal transformations at the cerebellum input stage. In a first hypothesis, plasticity was simply thought to weight Golgi cell inhibition of granule cells without specifying the synaptic process involved (Schweighofer et al., 2001). A more elaborated model implied the existence of LTP and LTD at the mossy fiber - Golgi cell synapses and anticipated their correlation with low-frequency oscillations of granular layer activity (Garrido et al., 2016). This work actually shows that Golgi cell plasticity is strictly bound to its electrophysiological properties and sets constraints to future models of the granular layer, that might allow to better understand the mechanisms of cerebellar learning and memory (Marr, 1969; Ito, 2008; Koziol et al., 2014; Sokolov et al., 2017; D’Angelo, 2019).

## Acknowledgments

This project has received funding from: the Human Brain Project (European Union’s Horizon 2020 Framework Program for Research and Innovation) under Grant Agreement No. 720270 (SGA1) and No. 785907 (SGA2) to ED; Centro Fermi grant MNL to ED. TS was supported by MNL, ST and FL by Human Brain Project. Special thanks to Javier DeFelipe (Department of Neuroanatomy and Cell Biology, Instituto Cajal (CSIC), Madrid, Spain) for Golgi cell morphological reconstruction.

